# Continuous Prediction of Leg Kinematics during Walking using Inertial Sensors, Smart Glasses, and Embedded Computing

**DOI:** 10.1101/2023.02.10.528052

**Authors:** Oleksii Tsepa, Roman Burakov, Brokoslaw Laschowski, Alex Mihailidis

## Abstract

Unlike traditional hierarchical controllers for robotic leg prostheses and exoskeletons, continuous systems could allow persons with mobility impairments to walk more naturally in real-world environments without requiring high-level switching between locomotion modes. To support these next-generation controllers, we developed a new system called *KIFNet* (Kinematics and Image Fusing Network) that uses lightweight and efficient deep learning models to continuously predict the leg kinematics during walking. We tested different sensor fusion methods to combine kinematics data from inertial sensors and computer vision data from smart glasses and found that adaptive instance normalization achieved the lowest RMSE predictions for knee and ankle joint kinematics. We also deployed our model on an embedded device. Without inference optimization, our model was 20 times faster than the previous state-of-the-art and achieved 20% higher prediction accuracies, and during some locomotor activities like stair descent, decreased RMSE up to 300%. With inference optimization, our best model achieved 125 FPS on an NVIDIA Jetson Nano. These results demonstrate the potential to build fast and accurate deep learning models for continuous prediction of leg kinematics during walking based on sensor fusion and embedded computing, therein providing a foundation for real-time continuous controllers for robotic leg prostheses and exoskeletons.

## I. Introduction

High-level control of robotic leg prostheses and exoskeletons is an important and challenging task. Most systems use a hierarchical architecture [1] where higher levels rely on automated systems to switch between different locomotion modes, each of which uses a separate controller with tuned parameters, and lower levels progress through the gait cycle while communicating with the robotic actuators. Recent studies [2]-[5] on continuous control have shown that it is possible to forward regress the leg kinematics from previous motions without the need to use high-level switching logic and separate controllers for different environments and/or gait phases. These systems have the potential of delivering more natural movements and higher flexibility in unstructured environments.

Computer vision systems like smart glasses can inform about the walking environment and sense obstacles before they are encountered, which other technologies like inertial measurement units (IMUs) are not fully capable of. Recent research [6]-[7] has shown that adding computer vision data can result in 7.9% and 7.0% improvement in root mean squared error (RMSE) of knee and ankle joint angle predictions, respectively, compared to using only inertial sensors; this was possible by pairing the inertial data with optical flow [8] mappings from the smart glasses. However, due to the high computational requirements of the proposed long short-term memory (LSTM)-based networks, their model was not capable of running real-time inference on an embedded system.

Motivated to translate these continuous prediction models to real-world applications with wearable robotics, we developed a new system called *KIFNet* (Kinematics and Image Fusing Network) to continuously predict the leg kinematics during walking in real-time with high accuracy on an embedded device. We studied different approaches to encoding inertial and computer vision data and evaluated different sensor fusion methods to combine the two representations. We also deployed our prediction model on an embedded device with the use of inference optimization to demonstrate the potential for continuous real-time control of robotic leg prostheses and exoskeletons. The code used to train and test our models is publicly available at https://github.com/Anvilondre/kifnet.

## II. Methods

### A. Multimodal Dataset

We used the multimodal dataset by [6] that includes fullbody kinematics from inertial sensors and RGB images with gaze from smart glasses. A detailed description is provided in [7]. Data was collected with 23 healthy young adults walking in 4 different annotated environments, including transitions, for a total of ∼12 hours. The environments included: classrooms with dense obstacle arrangements, an atrium with a sparse obstacle arrangement, and descending and ascending stairs. Subjects did not receive any walking instructions to preserve natural movement. Images and inertial data were collected at 30 Hz and 60 Hz, respectively. Kinematic data included 22 joint angles, each in 3 anatomical planes, from 17 inertial sensors. Our goal is to forward predict the right knee and ankle joint kinematics 1 second ahead of the user using only the lower-body sensors. Four input angles paired with the images were used (i.e., left knee, left ankle, left foot, and right hip). Inertial data was downsampled to 30 Hz to match the visual data. Images were downscaled to 320×240 resolution. We also omitted gaze from our study. The average gait cycle was ∼1 second.

We used the same training, validation, and testing splits as [6] such that out of the 23 subjects, 1 subject was used for testing, 3 subjects were used for validation, and the rest were used for training. Hyperparameter tuning and experiments were evaluated on the validation set and the testing set was only used for model predictions. We chronologically organized the data and split it into 7-second segments, resulting in 210 data points per segment. We calculated the RMSE over each segment and report the average and standard deviation across all segments. To measure inference speed on an embedded device, we computed the total time of *N* single-sample inferences and then divided it by *N* to calculate the average frames per second (FPS). We also report the FPS of the final model with inference optimization using TensorRT.

Whereas previous work [6] scaled the kinematics data from each trial separately from 0 to 1, we used normalization parameters from the entire training set to transform validation and testing sets. Although they reported scaled metrics, we reverted them to compare in the original scale. Since our approach required more time steps for prediction (45 instead of 10), we cropped the predictions by [6] by 35 frames for each of the 4 videos in the testing set. Overall, the testing dataset size decreased by 0.45%.

### B. Model Design and Training

To accurately the predict leg kinematics during walking, it is important to have a strong and robust representation of the user’s current state. In deep time-series analysis, the most common approaches encode the current state as a hidden state in recurrent neural networks (RNNs) or as a vector of concatenated previous states. We decided to use a standard feedforward neural network since RNNs tend to have significant computational requirements and poor scalability to longer sequences.

We used 46 previous time steps to forward regress the leg kinematics. Combined with 4 input joint angles, a total of 184 input values were passed to the feedforward network as an encoder. The output embedding dimension is a hyperparameter that was optimized together with the dimension of the image representation. The kinematic embeddings represent the current state of the user, and the image embeddings inform the model about the walking environment. We paired the current image from the smart glasses with the preceding 46 inertial measurements. We tested different computer vision backbones as an encoder, including the smallest version of MobileOne [9], MobileOne-S0, and MobileNetV2 [10]. Only the last stage (comprising ∼27.5% of trainable parameters) of MobileOne-S0 was optimized, keeping the rest of the model frozen. Similarly, the last 4 layers (comprising ∼10% of parameters) of MobileNetV2 were fine-tuned.

Models with a single decoder network tended to learn only one target joint well while compromising the other. Using separate decoders for the knee and ankle joints provided significantly more reliable results on both targets and increased the training stability. For the final model architecture (Fig. 1), we used MobileOne-S0 as the image encoder and a multi-layer perceptron (MLP) as a kinematic encoder. Both embeddings were passed to the sensor fusion module. After the fused embedding was computed, we passed it to two identical decoder networks but with different parameters.

**Figure 1.**
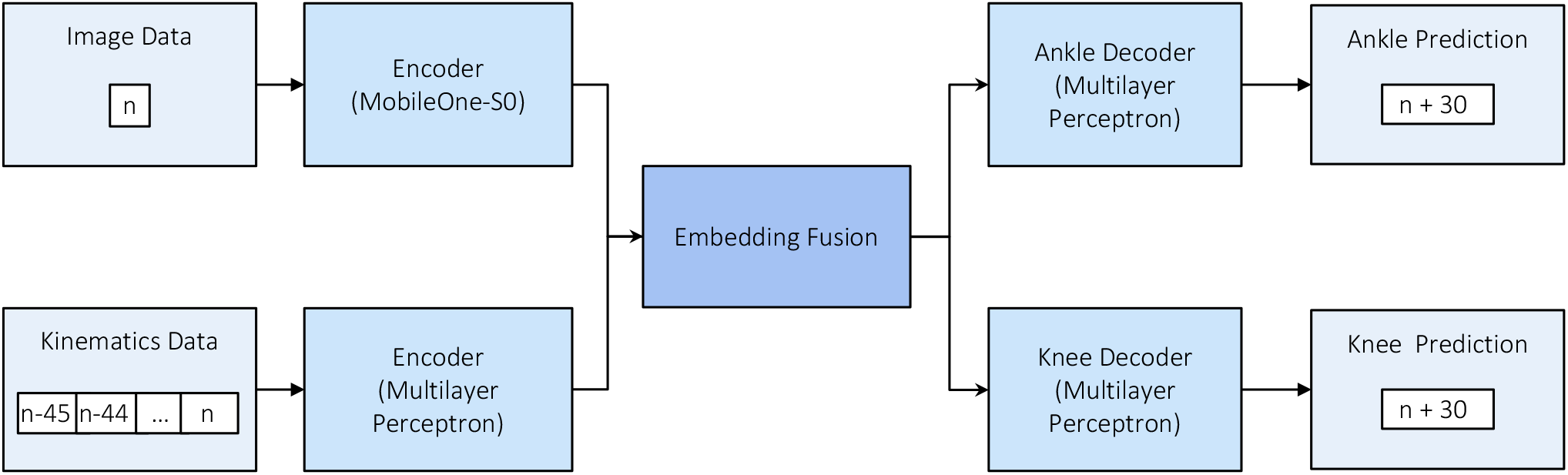
General model architecture. Kinematics data is represented as a vector of concatenated previous 46 measurements across 4 input joint angles. Computer vision data is represented as the last image from smart glasses. Both data modalities are passed to separate encoders and fused in the embedding fusion module. The resulting fusion vector is passed to two decoders for knee and ankle joint angle predictions, that are architecturally identical but have different parameters.

Training was performed on a single NVIDIA RTX A6000 and took ∼6.5 minutes per epoch. We used the Adam [11] optimizer and RMSE loss during training. The loss was computed by *L*_*total*_= 0.9 · *L*_*ankle*_ + 0.1 · *L*_*knee*_, where 0.9 and 0.1 are hand-selected coefficients that made training more stable and improved RMSE on both targets. The model was trained until the validation score did not improve for 5 epochs. To reduce overfitting, we added random Gaussian noise to the input training kinematics as a regularization method. After experimenting with batch normalization [12], layer normalization [13], dropouts [14], and L2 regularization [15], we found that neither technique worked particularly well for our application. Xavier [16] and He normal [17] weight initialization methods were tested but both increased overfitting. We also found that LeakyReLU [18] outperformed ReLU, ELU [19], and periodic activation functions [20].

### C. Sensor Fusion

Fusion of image and kinematic representations is challenging because the embeddings come from different latent spaces, making them difficult to combine in a manner that can be efficiently decoded into the final prediction. We evaluated different sensor fusion methods. Table 1 depicts the hyperparameters found by an automated search and later manually refined with systematic trial and error. Once our final model was developed, we iterated over each fusion method using this set of hyperparameters and measured the performance for comparison (Table 2). Note that the sizes of the fused embeddings differ such that the concatenation-based embeddings were twice as large as the others.

**Table 1.** Design and training hyperparameters for our final model.

| Hyperparameter | Value |
| --- | --- |
| Learning Rate | 1e-4 |
| Batch Size | 256 |
| Early Stopping (rounds) | 5 |
| Gaussian Noise (sigma) | 0.015 |
| Kinematics Encoder layers | [256, 256, 128] |
| Image Encoder | MobileOne-S0 (128 output) |
| Decoder layers | [64, 32] |
| LeakyReLU slope | -0.01 |

**Table 2.** Comparison of prediction accuracy for different data fusion methods on the testing set. Accuracy is represented by the RMSE of knee and ankle joint angle (°) predictions over the sliding windows of 7 seconds. Top-3 lowest RMSE for each joint are bolded.

| Fusion Method | Knee RMSE (°) | Ankle RMSE (°) |
| --- | --- | --- |
| Average | **7.30 ± 2.77** | 10.98 ± 4.35 |
| Maximum | 7.90 ± 2.51 | 9.87 ± 4.74 |
| Concatenation | 7.85 ± 2.59 | 9.85 ± 4.4 |
| Concat. + BatchNorm | 8.05 ± 2.98 | **9.47 ± 4.55** |
| Concat. + LayerNorm | **7.47 ± 2.75** | **9.78 ± 4.39** |
| Concat. + KL Loss | 7.57 ± 2.90 | 10.15 ± 4.32 |
| Dot Attention | 7.65 ± 2.87 | 11.93 ± 4.67 |
| MLP Attention | 7.51 ± 2.56 | 11.52 ± 4.78 |
| AdaIN (Kinematics-CV) | **6.85 ± 3.22** | **9.67 ± 4.72** |
| AdaIN (CV-Kinematics) | 8.32 ± 3.15 | 11.48 ± 4.73 |

To evaluate more complex aggregations, a baseline approach is needed. For these purposes, we first constructed fusion embeddings as an element-wise mean and maximum of both representations. A common data fusion method is to concatenate the embedding vectors. However, this introduces inconsistency in the latent space of the vector, which may lead to overfitting. To reduce this effect, 3 methods of latent space transformation were tested, including batch normalization, layer normalization, and Kullback–Leibler (KL) divergence [21], which pulls the vector distribution closer to normal:

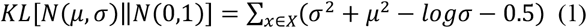

Another type of embedding aggregation, commonly used in natural language processing and graph neural networks, is based on the concept of Attention [22]. Fusion is achieved by dynamically computing the weighted sum of the embeddings, allowing the network to balance the representations depending on the input. Since we only have two embeddings, finding a single weight value is sufficient. We evaluated two methods of assigning weights, using a feedforward neural network (Fig. 2.a) and by calculating the dot product between the two vectors (Fig. 2.b); both of which followed by a Sigmoid activation function.

**Figure 2a.**
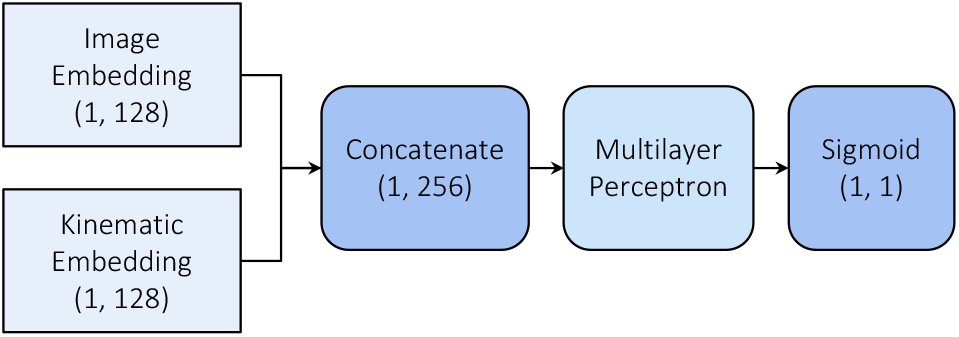
Calculating embedding weight using multilayer perceptron (MLP) attention with hidden dimensions of 64 and 32 with LeakyReLU activation between layers. We computed the sigmoid activation over the MLP output to get the final embedding weight.

**Figure 2b.**
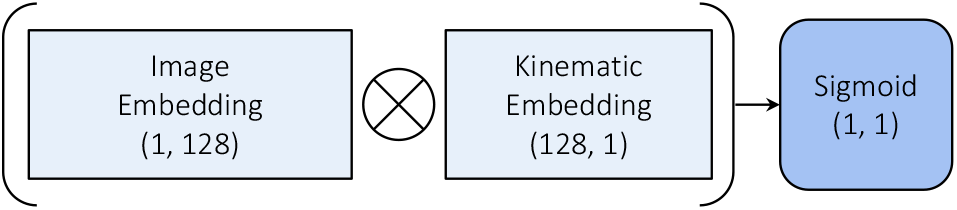
Calculating embedding weight using dot product attention. We computed the dot product between kinematics and transposed image embeddings followed by a sigmoid activation to get the final embedding weight.

We also tested adaptive instance normalization (AdaIN) [23], which takes content and style vectors as input and aligns the mean and variance of the content vector to match the style vector. Unlike batch normalization, instance normalization [24], or conditional instance normalization [25], AdaIN has no learnable parameters. Instead, it adaptively computes the affine parameters from the style input:

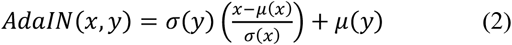

For the kinematic and image representations, one is selected as the content vector and another as the style vector. Because this operation is not commutative, both variants were tested.

### D. Embedded Deployment

We evaluated the inference speeds of our deep learning models on an NVIDIA Jetson Nano 2GB, which provides a reasonable trade-off between size, cost, and computational capacity. These embedded devices have the potential to support real-time control applications with robotic prosthetic legs and exoskeletons. We also did inference optimization using the TensorRT framework for our final models to approximate the potential inference speed in production. TensorRT can support different methods for inference optimization, including precision reduction, layer fusion, and efficient memory reuse. We used Python 3.8.13, TensorRT 8.0.1.6, and CUDA 11.3.

## III. Results

We found that space transformation methods such as batch normalization and layer normalization decreased RMSE compared to concatenated embedding, except for a slight increase in knee joint RMSE using batch normalization (Table 2). Adding Kulback-Leibler divergence did not improve performance, bringing visible underfitting. The averaging method worked well for the knee joint predictions but resulted in a high error for the ankle joint predictions. Attention-based aggregations performed poorly on the ankle. Adaptive instance normalization (kinematics as a content vector) achieved the best results with the lowest knee joint RMSE and second lowest ankle RMSE.

Without inference optimization, our best model was ∼20 times faster than the previous state-of-the-art [6] (Table 3), mainly due to having a faster backbone (MobileOne-S0 vs. PWC-Net) and using only one image for each prediction rather than sequences of 10. TensorRT was able to further speed up the MobileNetV2 and MobileOne-S0 models by 25 times and 23 times respectively. We reached over 125 FPS with the MobileOne model, which is more than sufficient for real-time prediction and control of wearable robotics.

**Table 3.** Comparison of inference speed and overall prediction accuracy between our models and the previous state-of-the-art [6]. Accuracy is represented by the RMSE of knee and ankle joint angle (°) predictions over the sliding windows of 7 seconds. Best values are bolded.

| Model | Frames per second (FPS) | Test Ankle RMSE ($^{\circ}$) | Test Knee RMSE ($^{\circ}$) |
| --- | --- | --- | --- |
| Previous Research [6] (PWC-Net) | 0.28 | $8.58 \pm 4.06$ | $12.30 \pm 5.62$ |
| KIFNet (MobileNetV2) | 2.88 | $8.5 \pm 2.80$ | $9.8 \pm 4.87$ |
| KIFNet (MobileOne-S0) | 5.50 | **$6.85 \pm 3.22$** | **$9.67 \pm 4.72$** |
| KIFNet (MobileNetV2 + TensorRT) | 74.46 | $8.5 \pm 2.80$ | $9.8 \pm 4.87$ |
| KIFNet (MobileOne-S0 + TensorRT) | **125.59** | **$6.85 \pm 3.22$** | **$9.67 \pm 4.72$** |

Compared to [6], we achieved higher prediction accuracies (up to 3 times higher) across different walking environments and targets (Table 4). We achieved 20.2% and 21.4% lower RMSE for overall ankle and knee joint kinematic predictions, respectively. Our model was able to utilize better computer vision data for training, which made the model more robust in different environments. To show that both data modalities influence the prediction, we quantify the relative importance of computer vision and kinematics data by interpreting our models with integrated gradients [26] (Fig. 3). The importance was calculated for each sample and aggregated to the environments and targets. Computer vision data was more valuable for the ankle joint predictions than for the knee. Interestingly, the importance of sensor modality was not consistent across different walking environments. For example, computer vision was most important for ankle joint predictions in transitions to stair ascent (TSA) compared to other locomotor activities but had the least effect on the knee joint predictions during incline stair transitions.

**Table 4.** Comparison of prediction accuracy for each walking environment between our model (KIFNet) and the previous state-of-the-art [6]. Accuracy is represented by the RMSE of knee and ankle joint angle (°) predictions over the sliding windows of 7 seconds. Best values are bolded.

|  | Ankle RMSE |  | Knee RMSE |  |
| --- | --- | --- | --- | --- |
|  | KIFNet System | Previous Research [6] | KIFNet System | Previous Research [6] |
| Classroom | **$6.92 \pm 2.06$** | $7.11 \pm 2.79$ | $10.25 \pm 4.69$ | **$9.82 \pm 3.14$** |
| Atrium | $6.58 \pm 2.66$ | **$6.09 \pm 1.88$** | **$9.23 \pm 4.95$** | $9.81 \pm 2.19$ |
| Stair Up | **$9.63 \pm 8.08$** | $12.66 \pm 3.24$ | **$9.63 \pm 2.76$** | $19.16 \pm 6.13$ |
| Stair Down | **$5.10 \pm 0.79$** | $18.69 \pm 2.68$ | **$6.62 \pm 0.93$** | $26.63 \pm 2.37$ |
| Door to Atrium | **$7.44 \pm 2.41$** | $9.26 \pm 3.11$ | $14.33 \pm 5.62$ | **$12.15 \pm 2.32$** |
| To Stair Ascent | **$6.54 \pm 1.24$** | $16.69 \pm 0.00$ | **$9.10 \pm 4.06$** | $22.70 \pm 0.00$ |
| Around Stairwell | **$5.57 \pm 0.97$** | $7.86 \pm 2.34$ | **$7.55 \pm 2.37$** | $9.14 \pm 2.10$ |
| To Stair Descent | **$4.68 \pm 1.05$** | $15.01 \pm 2.27$ | **$6.32 \pm 0.31$** | $21.10 \pm 0.57$ |
| To Atrium | **$5.00 \pm 0.96$** | $10.04 \pm 1.69$ | **$6.35 \pm 1.20$** | $19.17 \pm 6.64$ |
| Door to Classroom | **$7.50 \pm 3.96$** | $10.45 \pm 1.20$ | **$11.35 \pm 4.97$** | $15.50 \pm 2.96$ |
| Overall | **$6.85 \pm 3.22$** | $8.58 \pm 4.06$ | **$9.67 \pm 4.72$** | $12.30 \pm 5.62$ |

**Figure 3.**
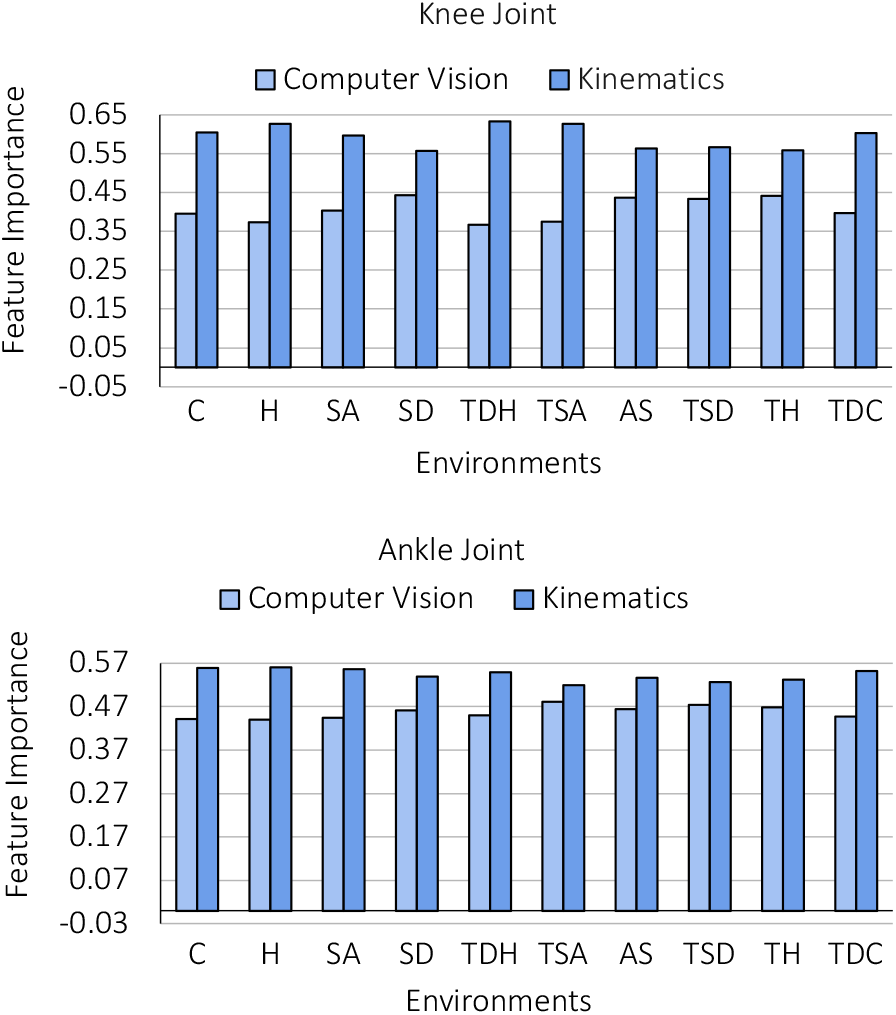
Comparing the importance of kinematics and computer vision inputs using integrated gradients [26]. C is classroom, H is atrium, SA is stair ascent, SD is stair descent, TSA is transitions to stair ascent, TDH is transition from door to atrium, AS is transitions around stairwell, TSD is transitions to stair descent, TH is transitions to atrium, and TDC is transitions from door to classroom.

## IV. Discussion

In this study, we developed a new deep learning-based system (*KIFNet*) to forward predict the leg kinematics during walking for continuous control of robotic leg prostheses and exoskeletons. We used computer vision data from smart glasses and kinematics data from inertial sensors and provided technical contributions in the areas of multimodal sensor fusion and lightweight and efficient deep learning for real-time inference on embedded devices. Our system builds on the latest advancements in neural network design for mobile vision applications [9]-[10], which balances computational cost with prediction accuracy. We also compared different embedding fusion techniques and are one of the first systems to use adaptive instance normalization [23] to integrate time-series and computer vision data for regression problems. Note that, whereas previous applications of computer vision in wearable robotics have focused on classification [27], here we focused on regression to design continuous controllers.

Without inference optimization, our model was ∼20 times faster than the previous state-of-the-art [6] due to having a faster backbone (MobileOne-S0 vs. PWC-Net) and using only one image for each prediction rather than sequences of 10. We also achieved higher prediction accuracies (up to 3 times higher) across different walking environments and targets. Overall, we achieved ∼20% lower RMSE for ankle and knee joint kinematic predictions, respectively. With TensorRT inference optimization, our MobileOne model reached 125 FPS on an NVIDIA Jetson Nano, which is an order of magnitude greater than what is needed for real-time prediction and control of wearable robotics. TensorRT was able to further speed up our MobileNetV2 and MobileOne-S0 models by 25 times and 23 times, respectively.

Despite these developments, our study has limitations. For example, although our deep learning model was deployed on an embedded device and showed real-time inference, our predictions were not directly integrated to robot control hardware, which is eventually needed for proof of concept. Future work is needed in systems integration in order to evaluate the feasibility of using *KIFNet* for continuous control of robotic leg prostheses and exoskeletons. These continuous systems, compared to traditional hierarchical controllers, have the potential to allow persons with mobility impairments to walk more naturally in real-world environments without requiring high-level switching between locomotion modes.

## Acknowledgment

We thank Abhishek Sharma and Erick Rombokas from the University of Washington for their assistance with the multimodal dataset. This research is dedicated to the people of Ukraine in response to the 2022 Russian invasion.

## Notes

* O. Tsepa and R. Burakov provided equal contribution, listed randomly. Research supported by the AGE-WELL Networks of Centres of Excellence program, the Department of Computer Science, University of Toronto, and the Vector Institute for Artificial Intelligence.

### Competing Interest Statement

The authors have declared no competing interest.

